# Identification of a novel papillomavirus from a New Zealand fur seal (*Arctocephalus forsteri*) with oral papilloma-like lesions

**DOI:** 10.1101/2023.11.20.567757

**Authors:** Jonathon C.O. Mifsud, Jane Hall, Kate Van Brussel, Karrie Rose, Rhys H. Parry, Edward C. Holmes, Erin Harvey

**Affiliations:** Sydney Institute for Infectious Diseases, School of Medical Sciences, The University of Sydney, Sydney, NSW 2006, Australia; Centre for Planetary Health and Food Security, School of Environment and Science, Griffith University, Nathan, Queensland, Australia; Australian Registry of Wildlife Health, Taronga Conservation Society Australia, Mosman, NSW 2088, Australia; School of Chemistry and Molecular Biosciences, The University of Queensland, Brisbane, QLD 4067, Australia

**Keywords:** *Papillomavirus*, *Taupapillomavirus*, *Herpesvirus*, New Zealand fur seal, pinniped, *Arctocephalus forsteri*

## Abstract

Despite being the predominant seal species in the Australian-New Zealand region and serving as a key indicator of marine environmental health, little is known about infectious diseases in New Zealand fur seals (Long-nosed fur seal; *Arctocephalus forsteri*). Several papillomaviruses have been identified in earless seals and sea lions, with the latter linked to cutaneous plaques and invasive squamous cell carcinoma. To date, no papillomaviruses have been reported in fur seals. We used traditional veterinary diagnostic techniques and metatranscriptomic sequencing of tissue samples to investigate the virome of New Zealand fur seals. We identified a novel papillomavirus, provisionally termed Arctocephalus forsteri papillomavirus 1 (AfPV1) in an animal with clinically and histologically identified oral papilloma-like lesions. RT-PCR confirmed the presence of AfPV1 only in oral papilloma samples from the affected individual. Phylogenetic analysis of the complete 7,926 bp genome of AfPV1 revealed that it clustered with taupapillomaviruses found in related Carnivora species. In addition, we identified the partial genome of a novel Gammaherpesvirus, Arctocephalus forsteri gammaherpesvirus 1 (AfGHV1), in a different individual without pathological evidence of viral infection. These findings highlight the need for further research into the disease associations and impact of undiagnosed and novel viruses on New Zealand fur seals.

## 1. Introduction

New Zealand fur seals (Long-nosed fur seal; *Arctocephalus forsteri*) are the most abundant seal species in the Australian-New Zealand region (Goldsworthy et al., 2003). Their breeding colonies span the southern coast of Australia, ranging from Western Australia to Tasmania, as well as the coastlines and offshore islands of New Zealand and its subantarctic islands (Shaughnessy, 1999). As long-lived marine mammals that feed at high trophic levels, New Zealand fur seals serve as important marine sentinels and offer insights on the health of aquatic ecosystems (Bossart, 2011).

Despite their importance, our understanding of infectious diseases in New Zealand fur seals, particularly those due to viruses, remains limited. A single circovirus has been identified from a New Zealand fur seal faecal sample (Sikorski et al., 2013), although no details on the health status of the seal or the potential for pathogenicity were provided. The impact viruses can have on other seal species is well established, with numerous viral diseases described, including urogenital carcinoma associated with Otarine herpesvirus 1 in California sea lions (*Zalophus californianus*) and a South American fur seal (*A. australis*) (Dagleish et al., 2013; Deming et al., 2021) and vesicular exanthema associated with San Miguel sea lion virus reported in sea lions, fur seals, and elephant seals along the western coast of the United States (Bossart and Duignan, 2019). Viral diseases in seals can have far-reaching consequences (Colegrove et al., 2005). For instance, outbreaks of seal influenza A (H10N7) across Europe in 2014 (Bodewes et al., 2015) and highly pathogenic avian influenza A (H5N1) in New England, USA, in 2023, both caused mass mortalities of harbour seals (*Phoca vitulina*) (Puryear et al., 2023). Additionally, seals may have the potential to affect human health through zoonotic virus transmission, particularly during close interactions such as stranding and mass mortality events (Abdelwhab and Mettenleiter, 2023).

The *Papillomaviridae* are a family of non-enveloped, double-stranded DNA viruses, typically 7,500 bp in length, that exhibit high host and tissue specificity, primarily infecting skin and mucosal surfaces. Infection can manifest in a range of ways, from asymptomatic, to the formation of self-resolving papillary masses, to malignant epithelial cancers (Syrjänen, 2018). This family of viruses has a broad host range, spanning diverse species and environments. Apart from the genus *Alphapapillomavirus* within the subfamily *Secondpapillomavirinae,* which is known to infect fish, all other papillomaviruses belong to the subfamily *Firstpapillomavirinae* and are associated with reptiles, birds, and mammals, from both terrestrial and aquatic environments (Van Doorslaer et al., 2018).

Few papillomaviruses have been identified in pinnipeds. Zalophus californianus papillomavirus (genus *Dyonupapillomavirus*) was reported from California sea lions (*Zalophus californianus*) presenting with axillary and preputial papillomatous lesions (Rivera et al., 2012). This virus has been associated with sporadic cases of both *in situ* and invasive squamous cell carcinoma (Luff et al., 2018). In addition, various genera of papillomaviruses have been identified in the faeces of apparently healthy Weddell seals (*Leptonychotes weddellii*) (Smeele et al., 2018).

Given the potential for pinniped viruses to cause population-level effects, and the risk posed to veterinarians and rehabilitators from zoonotic diseases, it is crucial to expand our knowledge of seal viruses. Accordingly, we employed traditional veterinary diagnostic techniques in conjunction with metatranscriptomic sequencing of a variety of tissues from seals, with or without clinical or pathological signs of viral infection, to explore the virome of New Zealand fur seals.

## 2. Methods

### 2.1 Sample collection and processing

Samples from live seals were collected by a veterinarian between 2003 and 2021 for diagnostic purposes under a License to Rehabilitate Injured, Sick or Orphaned Protected Wildlife (no. MWL000100542) issued by the New South Wales (NSW) Department of the Environment. Samples from deceased, beach cast, or euthanased seals were collected by the Australian Registry of Wildlife Health (Registry), a conservation science program of Taronga Conservation Society Australia, during routine necropsy for disease surveillance in accordance with NSW National Parks and Wildlife Act 1974, section 132c, Scientific Licence number SL100104. Tissue samples comprising various skin and mucosa samples, liver, brain, lung, and kidney were variably collected from 18 New Zealand fur seals and stored at –80 °C until RNA extraction. A set of tissues representing each organ system was also collected into 10% neutral buffered formalin, embedded in paraffin wax, sectioned and mounted on a glass slide, stained with hematoxylin and eosin, and examined by light microscopy at 200, 400, and 1000x magnification.

### 2.2 RNA extraction and metatranscriptomic sequencing

Individual tissue aliquots were placed into 600 μl of lysis buffer containing 0.5% foaming reagent (Reagent DX, Qiagen) and 1% β-mercaptoethanol (Sigma-Aldrich), and tissue was homogenised using a TissueRuptor (Qiagen) at a speed of 5,000 rpm for up to one minute. The homogenate was centrifuged at maximum speed (15,200 rpm) for three minutes to eliminate any remaining tissue residue. RNA was then extracted from the supernatant using the RNeasy Plus Mini Kit (Qiagen), following the manufacturer’s protocol. Extracted RNA was combined by tissue type into 14 pools with a median of three samples per pool (minimum = 1, maximum = 5) (Supplementary Table 1). Sequencing libraries were constructed using the TruSeq Total RNA Library Preparation Protocol (Illumina). Host ribosomal RNA (rRNA) was depleted using the Ribo-Zero Plus Kit (Illumina) and paired-end sequencing (150 bp) was performed on the NovaSeq 6000 platform (Illumina). Library construction and sequencing were performed by the Australian Genome Research Facility (AGRF).

### 2.3 Identification of novel virus sequences

Virus identification followed the BatchArtemisSRAMiner pipeline (Mifsud, 2023). Briefly, sequencing reads underwent quality trimming and adapter removal using Trimmomatic (v0.38) with parameters SLIDINGWINDOW:4:5, LEADING:5, TRAILING:5, and MINLEN:25, prior to assembly (Bolger et al., 2014). *De novo* assembly was conducted using MEGAHIT (v1.2.9) (Li et al., 2015). Assembled contigs were compared to the RdRp-scan RdRp core protein sequence database (v0.90) (Charon et al., 2022) and the protein version of the Reference Viral Databases (v23.0) (Goodacre et al., 2018) using DIAMOND BLASTx (v2.0.9) with an E-value cut-off of 1 × 10^−5^ (Buchfink et al., 2021). To exclude potential false positives, contigs with hits to virus sequences were used as a query against the NCBI nucleotide database (as of March, 2022) using BLASTn (Camacho et al., 2009) and the NCBI non-redundant protein (nr) database (as of March, 2022) using DIAMOND BLASTx. Using BLASTx and BLASTn matches, virus-like contigs associated with non-vertebrate hosts were excluded.

### 2.4 Arctocephalus forsteri papillomavirus 1 RT-PCR

RT-PCR was performed on total RNA from the five individual samples that made up library SL16 using primers designed based upon the novel Arctocephalus forsteri papillomavirus 1 (AfPV1) fragments obtained by metatranscriptomic sequencing. SuperScript IV One Step RT-PCR (Invitrogen) and the forward primer 5’ TGGAACGTTGACCTGAGAGA 3’ and reverse primer 5’ AAGGATACGGTCCGTTCTGA 3’ were used to amplify a missing 689 bp section between the L1 and E6 genes. A second set of primers, forward primer 5’ ATACACTCCGTCTTGGGACG 3’ and reverse primer 5’ CAGTTACAAAGCTTCGAGGGT 3’, was used to check the region surrounding the stop codon of E2. The resulting amplicon product was subsequently used for Sanger sequencing at the AGRF.

### 2.5 Arctocephalus forsteri gammaherpesvirus 1 sequencing

A previously published viral particle enrichment protocol (Conceição-Neto et al., 2015) was used to extract and randomly amplify nucleic acids from the oral tissue of a juvenile male seal TARZ-10741. Tissue was homogenised, filtered, centrifuged, and nuclease treated prior to viral nucleic acid extraction with the QIAamp viral RNA mini kit (Qiagen) (Chong et al., 2019; Conceição-Neto et al., 2015). Following this, nucleic acids were randomly amplified using the Whole Transcriptome Amplification kit (WTA2, Sigma Aldrich) with modifications (Conceição-Neto et al., 2015), and purified using the GenElute PCR cleanup kit (Sigma Aldrich). The DNA library was prepared using the Illumina DNA M preparation kit and sequenced on the Illumina NovaSeq 6000 platform at the AGRF. As the standard clean-up protocol depletes amplicons < 500 bp, these amplicons underwent Illumina purification using beads at a ratio of 1.8x volume to supernatant.

### 2.6 Genome analysis and annotation

To examine the genome coverage of each virus, sequence reads were mapped onto virus-like contigs using BBMap (v37.98) (Bushnell, 2014), and areas of heterogeneous coverage were manually checked using Geneious (v11.0.9). Where possible, the extremities of contigs were manually extended and re-submitted to read mapping until the contig appeared complete or no overhanging extremities were observed. Sequences of vector origin were detected using VecScreen (https://www.ncbi.nlm.nih.gov/tools/vecscreen/) and removed. GetORF from EMBOSS (v6.6.0) was used to predict open reading frames (ORFs) (Rice et al., 2000). To annotate protein functional domains, the InterProScan software package (v5.56) was used with the TIGRFAMs (v15.0), SFLD (v4.0), PANTHER (v15.0), SuperFamily (v1.75), PROSITE (v2022_01), CDD (v3.18), Pfam (v34.0), Hamap (v 2023_01), SMART (v7.1), PRINTS (v42.0), PIRSF (v3.10) and CATH-Gene3D databases (v4.3.0) (Jones et al., 2014). The completeness and quality of viral sequences were assessed by visual inspection and the CheckV pipeline (Nayfach et al., 2021). AfPV1 gene expression levels were calculated using htseq-count (v2.0.3) with non-default parameters “-s reverse –-nonunique fraction”. AfPV1 binding sites were predicted by manual sequence comparisons, while GC content was calculated in Geneious with a sliding window of 40 nucleotides. AfPV1 spliced ORFs E1åE4 and E8åE2 were manually predicted by aligning AfPV1 to Canis familiaris papillomavirus 19 isolate tvmb1 (KX599536). Support for the splice junctions was assessed using ViReMa (v0.25) (Sotcheff et al., 2023). Circos (v0.69-6) was used to produce the circular genome graphs for AfPV1 (Krzywinski et al., 2009). The marker gene cytochrome c oxidase subunit I (COX1) was identified through querying the contig set against the nr database using DIAMOND BLASTx. The abundances of COX1 and viral transcripts (both AfPV1 and AfGHV1) were determined by individually mapping the SL16 reads to each using RNA-Seq by Expectation Maximization (RSEM) software (v1.3.0) (Li and Dewey, 2011).

### 2.7 Assessment of sequencing library composition

To identify any possible contaminant sequences or coinfecting bacteria or fungi, contigs from the SL16 library were aligned to the custom NCBI nt database using the KMA aligner and the CCMetagen program (Clausen et al., 2018; Marcelino et al., 2020). Species related to known pathogens of mammals were manually confirmed through BLASTn and read mapping against reference genes using BBMap.

### 2.8 Phylogenetic analysis

Phylogenetic trees of the putative papillomavirus and herpesvirus sequences identified here were inferred using a maximum likelihood approach. Representative genomes (n = 117) from each of the papillomavirus genera were downloaded from The Papillomavirus Episteme (PaVE) (https://pave.niaid.nih.gov/) (Van Doorslaer et al., 2017). The amino acid sequences of four genes (L1, L2, E1 and E2) were obtained for these sequences along with the novel papillomavirus identified here and individually aligned using MAFFT (v7.402) (Katoh and Standley, 2013), quality trimmed using trimAl (v1.2) (Capella-Gutiérrez et al., 2009), and concatenated to form a single alignment. All phylogenetic trees were estimated using IQ-TREE2 (Minh et al., 2020). Branch support was calculated using 1,000 bootstrap replicates with the UFBoot2 algorithm and an implementation of the SH-like approximate likelihood ratio test within IQ-TREE2 (Anisimova et al., 2011). The best-fit model of amino acid substitution was determined using the Akaike information criterion (AIC), the corrected AIC, and the Bayesian information criterion with the ModelFinder function in IQ-TREE2 (Kalyaanamoorthy et al., 2017). This process was then repeated with a subset of papillomaviruses, namely the taupapillomaviruses and the related gamma– and pipapillomaviruses.

### 2.9 Data availability

All *A. forsteri* sequence reads are available on the NCBI Sequence Read Archive (SRA) under BioProject PRJNA1013207. All viral genomes assembled in this study have been deposited in the GenBank and assigned accession numbers OR531434, OR590706 and OR590707. The sequences, alignments, phylogenetic trees generated in this study is available at https://github.com/JonathonMifsud/Identification-of-a-novel-papillomavirus-in-a-New-Zealand-Fur-seal-with-oral-papilloma

## 3. Results

### 3.1 Overview of metatranscriptomic data

In total, 14 metatranscriptomic libraries were constructed from pools of tissue from 18 individual New Zealand fur seals (library statistics are summarised in Supplementary Table 1). The extracted RNA was pooled according to tissue type. No mammalian-associated viruses were identified from the brain, liver, lung, or kidney. However, a novel papillomavirus and a novel herpesvirus were recovered from oral tissue library SL16 and are discussed in further detail below.

### 3.2 Identification of a novel papillomavirus in a seal with oral papilloma

In October 2002, an immature male New Zealand fur seal (Registry #3254) was taken into rehabilitation care after being found hauled out on a beach near Narooma, New South Wales (NSW), Australia, in an emaciated body condition and exhibiting dehydration, severe anaemia, and several deep skin wounds over the right hip and right hind flipper, presumably associated with a failed predation attempt by a shark. Upon examination, the seal was noted to have several small raised, sometimes pedunculated and coalescing papillary masses on the roof of the mouth with similar, but smaller lesions, evident on the caudoventral right and ventral left aspects of the tongue (Figure 1A,B). There was also a circumferential zone of mucosal pallor around the right mandibular canine tooth, which was not biopsied (Figure 1C).

**Figure 1.**
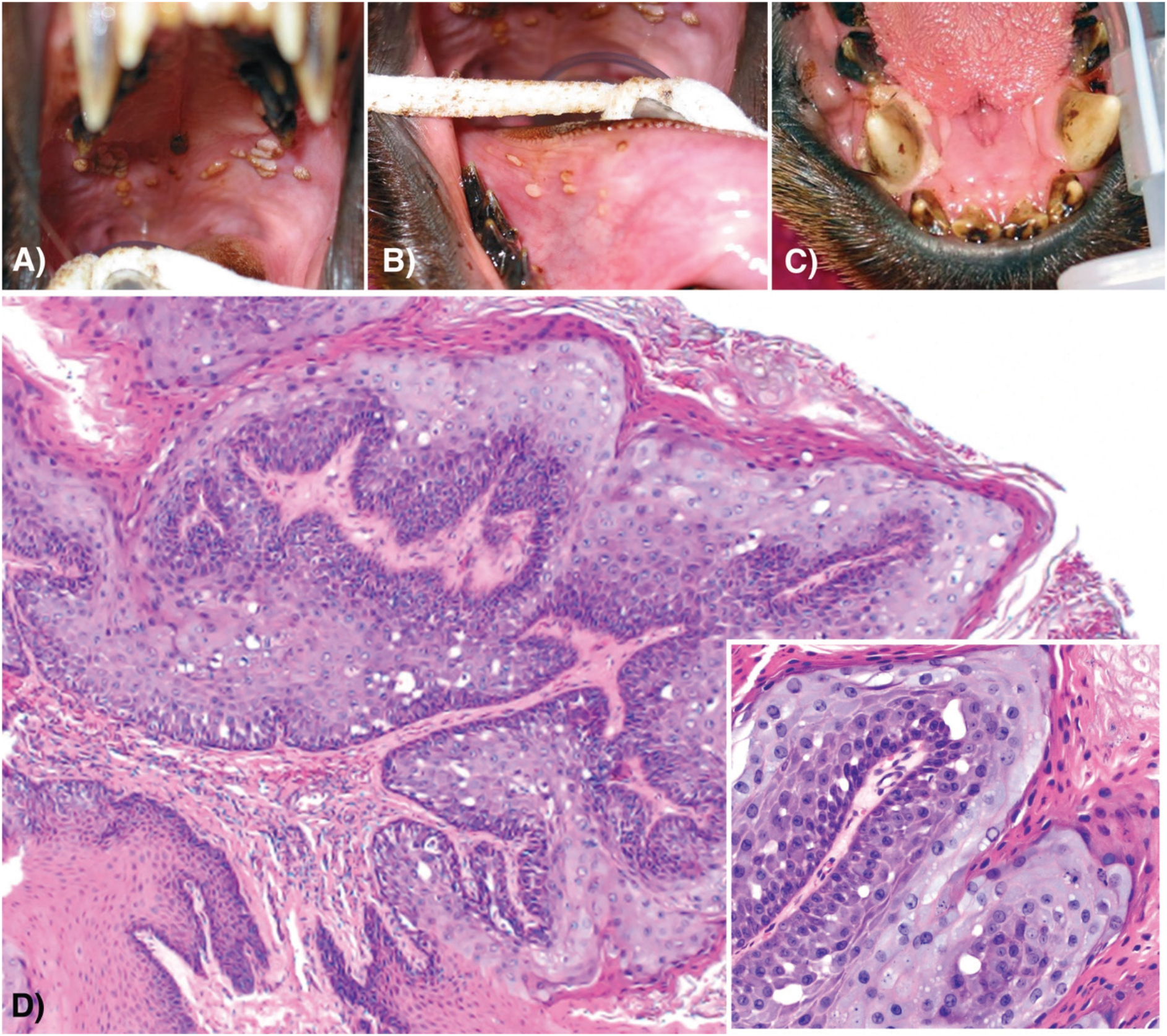
Multifocally coalescing sessile to papillary proliferations of the palatine and lingual epithelium of a New Zealand fur seal Registry #3254 (A, B). A zone of mucosal pallor surrounding the mandibular canine tooth (C). Demarcated lingual epithelial proliferation with basophilia and expansion of the stratum spinosum and stratum corneum (D – higher magnification of the lesion, inset).

Microscopic examination of multiple formalin-fixed oral lesion biopsies revealed sharply demarcated zones of lingual epithelial proliferation characterised by basophilia, binucleation of cells throughout the stratum basale, and an epithelium irregularly thickened by expansion of the stratum spinosum and stratum corneum (Figure 1D). Cells in the stratum spinosum were large and contained abundant basophilic cytoplasm. Moderate numbers of necrotic cells, with pyknotic or karyorrhectic nuclei, and intercellular oedema were multifocally evident throughout the affected stratum spinosum. The lingual lamina propria beneath the epithelial lesions appeared normal.

#### 3.2.1 Papillomavirus genome construction and confirmation

Several papillomavirus-like contigs were assembled from library SL16. Using a combination of read mapping and RT-PCR, a complete circular genome of 7,926 bp here termed Arctocephalus forsteri papillomavirus 1 (AfPV1) was recovered, which was confirmed by CheckV (100% genome completion, high confidence) (Figure 2). AfPV1 was present in high abundance in SL16 with 0.18% (n = 130,197 reads) of total library reads mapping to the genome, slightly less than the abundance of the mitochondrial COX1 gene in this library (0.27%, n = 195,218 reads). RT-PCR confirmed that AfPV1 was limited to seal Registry #3254, with both tissue samples taken from under the tongue testing positive. No other tissues or blood samples were available from this individual.

**Figure 2.**
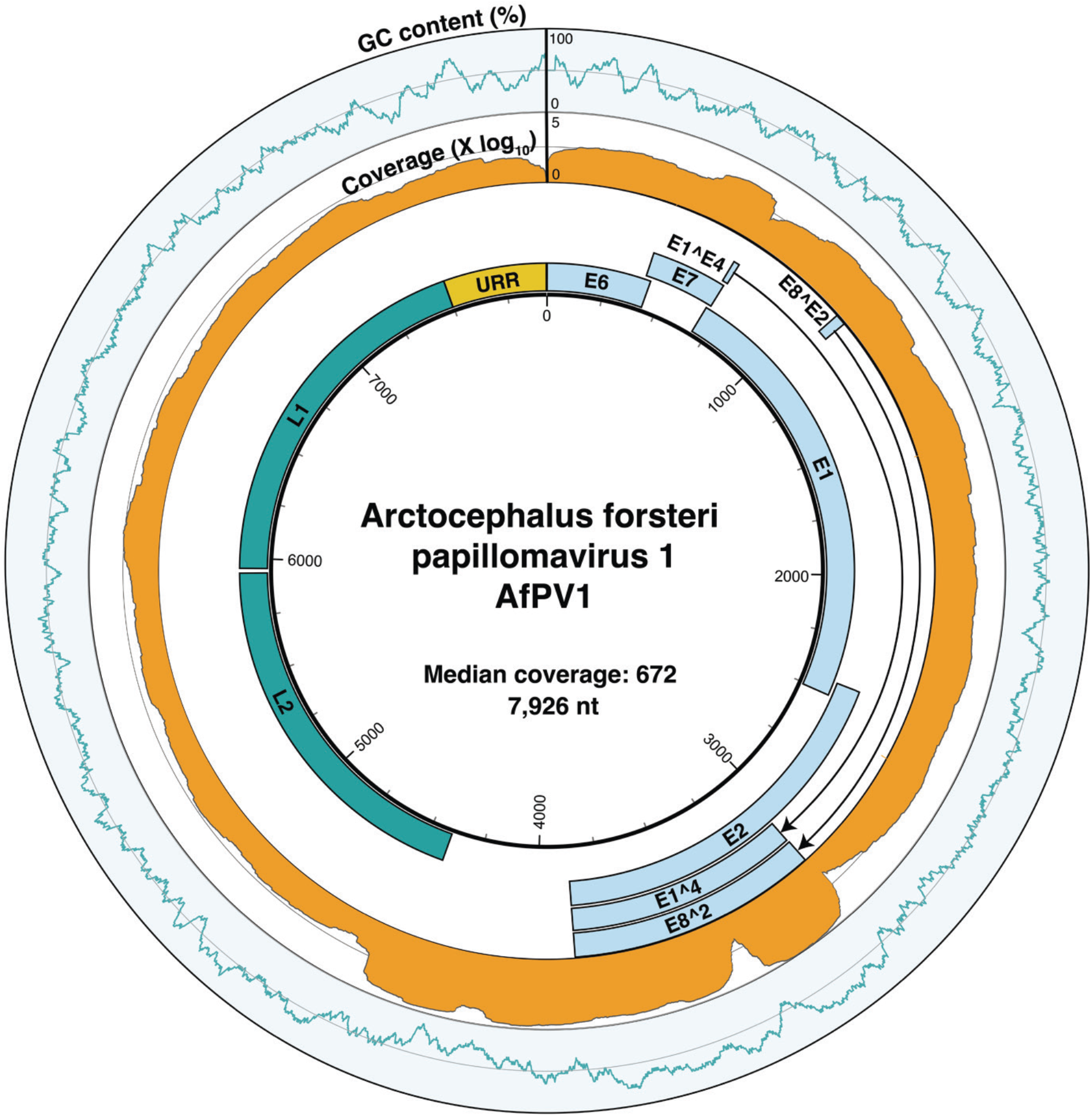
Genome organisation of Arctocephalus forsteri papillomavirus 1 (AfPV1). The outer ring indicates the percentage GC content (blue) over a 40bp sliding window, the middle ring represents read coverage across the genome, log_10_ transformed (orange), and the inner ring illustrates the predicted ORFs and the upstream regulatory region of AfPV1. The “A” in the first start codon of the ORF E6 is assigned as position one.

#### 3.2.2 Evolutionary relationship and genomic properties of AfPV1

AfPV1 falls as a distinct lineage within the papillomaviruses, with its closest relative Mustela putorius papillomavirus 1 (MpPV1) (NC_022253) isolated from a European polecat (*Mustela putorius*, family *Mustelidae*) with which it shares 66% nucleotide identity across the genome (Figure 3A). However, the degree of similarity between AfPV1 and MpPV1 varied between the six genes tested (E1, E2, E6, E7, L1 and L2), ranging from 50% (E2) to 74% (L1) nucleotide identity. To determine the evolutionary history of AfPV1 a phylogenetic analysis was conducted using a concatenated alignment of four genes (L1, L2, E1 and E2) as per the ICTV guidelines (Van Doorslaer et al., 2018). This analysis placed AfPV1 within the genus *Taupapillomavirus,* forming a clade with MpPV1 and a papillomavirus associated with the Weddell seal, Leptonychotes weddellii papillomavirus 5 (LwPV5), albeit with weak bootstrap support (SH-aLRT = 43% and UFboot = 64%) (Figure 3B). Given that the ICTV species demarcation (Van Doorslaer et al., 2018) for the taupapillomaviruses is <70% nucleotide identity across the genome, we suggest that AfPV1 represents a novel species within the genus *Taupapillomavirus* (family *Papillomaviridae*).

**Figure 3.**
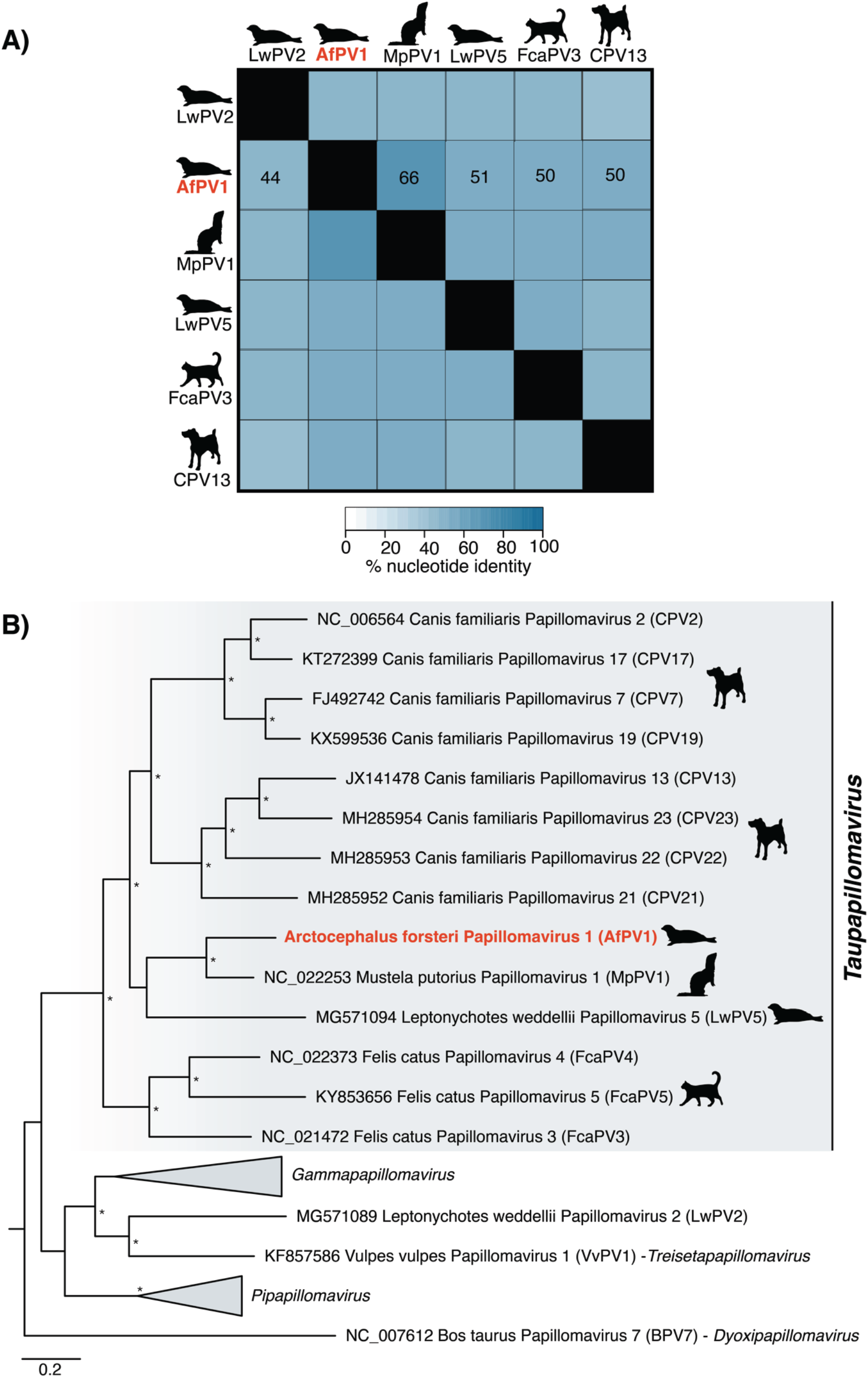
(A) Percent identity matrix of representative taupapillomaviruses for each host species compared to AfPV1. (B) Phylogenetic relationship of the taupapillomaviruses and other closely related genera. An ML phylogenetic tree based on the conserved amino acid sequences of the L1, L2, E1, E2 genes shows, in red, the topological position of Arctocephalus forsteri papillomavirus 1 (AfPV1), in the context of its closest relatives. Animal silhouettes depict the virus-host associations for the taupapillomaviruses. All branches are scaled to the number of amino acid substitutions (model LG+F+I+G4) per site, and the tree is midpoint rooted for clarity only. An asterisk indicates node support where SH-aLRT >= 80% and UFboot >= 95%.

The genome organisation of AfPV1 is consistent with other taupapillomaviruses, comprising eight open reading frames (ORFs) (E1, E2, E4, E6, E7, E8, L1 and L2) as well as the spliced ORFs of E1åE4 and E8åE2. The presence of the E8åE2 and E1åE4 splice junctions was supported by 89 and 13,700 reads, respectively. No E5 ORF was detected. Six of the eight ORFs had matches to domains associated with the E1, E2, E6, E7, L1 and L2 ORFs (Figure 2). No domains were detected for the E4 and E8 ORFs. A premature stop codon, which resulted in E2 being 238 bp (79 aa) shorter than its closest relatives, was present. This stop codon was in a region of high coverage and was confirmed by Sanger sequencing. Two zinc-binding sites (CXXC-X29-CXXC) were found in E6, although E6 lacked a PDZ-binding motif (ETQL) in its C-terminus. An alternative pRB-binding site (retinoblastoma tumour suppressor-binding domain) (LXSXE) was detected in E7, consistent with other taupapillomaviruses (Smeele et al., 2018; Wang et al., 2010). A cyclin RXL motif (KRRLF) and ATP-binding site (GXXXXGK[T/S]) were detected in E1. The upstream regulatory region (URR), defined as the region between the L1 stop codon and E6 start codon, was 430 bp in length. The URR contained four E2-binding sites (ACCN2-11GGT), one Nf1-binding site (TTGGC), one palindromic E1-binding site (ATTGTTXXXAACAAT) and a TATA box. The predicted ORFs, protein products, and binding sites are shown in Supplementary Table 2.

#### 3.2.3 Gene expression of AfPV1

Gene expression analysis of AfPV1 revealed variation among its genes (Supplementary Table 3). Notably, E2 displayed relatively high expression levels (194,773 reads per kilobase per million mapped reads [RPKM]). Expression was concentrated in specific regions, encompassing nucleotides 3,048-3,331 and 3,397-3,872, which corresponded to the predicted alternatively spliced isoforms, E8åE2 and E1åE4, each recording RPKM values of 315,227 and 315,461, respectively. In contrast, the remaining early genes E1, E6, and E7 exhibited comparatively lower RPKM values of 39,158, 12,894, and 10,751, respectively. The late gene L2 exhibited higher expression (63,078 RPKM) than L1 (4,693 RPKM), although this was predominantly observed at the beginning of the L2 gene (nucleotides 4,402-4,510).

#### 3.2.4 Detection of chimeric host AfPV1 contigs

A chimeric contig was assembled, containing the complete AfPV1 genome along with predicted *A. forsteri* genes (Supplementary Figure 1A). The circular virus genome appears to have been split at approximately the midpoint of the E2 gene (AfPV1 nucleotides 3,225-3,330). On the 5’ end of the E2 gene, a 918bp ORF was predicted, which was identical to the sea lion mitochondrial ribosomal protein L2 (MRPL2) gene (XM_027602888.1). Adjacent to the 3’ end E2 gene, an 824bp fragment was identified, exhibiting 99% sequence similarity with the sea lion purinergic receptor P2Y2 (P2RY2) gene (XM_028115895). When aligned with XM_028115895, this fragment was found downstream of the P2RY2 ORF (Supplementary Figure 1B, C). Read coverage is consistently >50X throughout the contig, although it drops to 7X and 41X at the predicted P2RY2-E2 and E2-MRPL2 junctions, respectively (Supplementary Figure 1B).

### 3.3 Identification of a novel seal gammaherpesvirus

Seal Registry #10741 was an immature male rescued at Cronulla, NSW, Australia, in September 2015. The seal presented in an emaciated body condition with various traumatic injuries, including open and infected metatarsal and phalangeal fractures, and was euthanased due to a poor prognosis for recovery. Gross and histopathological examinations confirmed the animal’s emaciated state, the presence of intestinal helminths (within the expected range for a free-ranging pinniped), severe traumatic injuries, and a neutrophilia consistent with a systemic inflammatory response secondary to infected wounds. Genital and oral mucosal tissue samples were taken from this individual for RNA extraction and pooled to form library SL16, the same library in which the papillomavirus contigs were identified.

In the SL16 library, several fragmented contigs exhibited similarities to gammaherpesviruses, comprising 0.005% of the total reads (n = 3,503). RT-PCR analysis confirmed that this sequence was exclusive to the oral tissue of seal Registry #10741, with no presence in the genital tissue of this individual. Two contigs containing a partial polymerase gene (3,460 bp) and the major capsid protein (12,493 bp) were assembled from the RNA libraries. DNA viral particle enrichment yielded related reads (6,310 reads when mapped to NC_035117, the closest relative with a 60% nucleotide identity cut-off), but we were unable to substantially extend the contigs recovered through RNA-seq or recover other core gammaherpesvirus genes. The partial polymerase sequence shared 72% amino acid identity with phocid herpesvirus 7 (AJA71665.1). Phylogenetic analysis revealed that this sequence grouped within a clade of pinniped-associated gammaherpesviruses within the genus *Rhadinovirus* (Figure 4). We tentatively assign this virus as Arctocephalus forsteri gammaherpesvirus 1 (AfGHV1).

**Figure 4.**
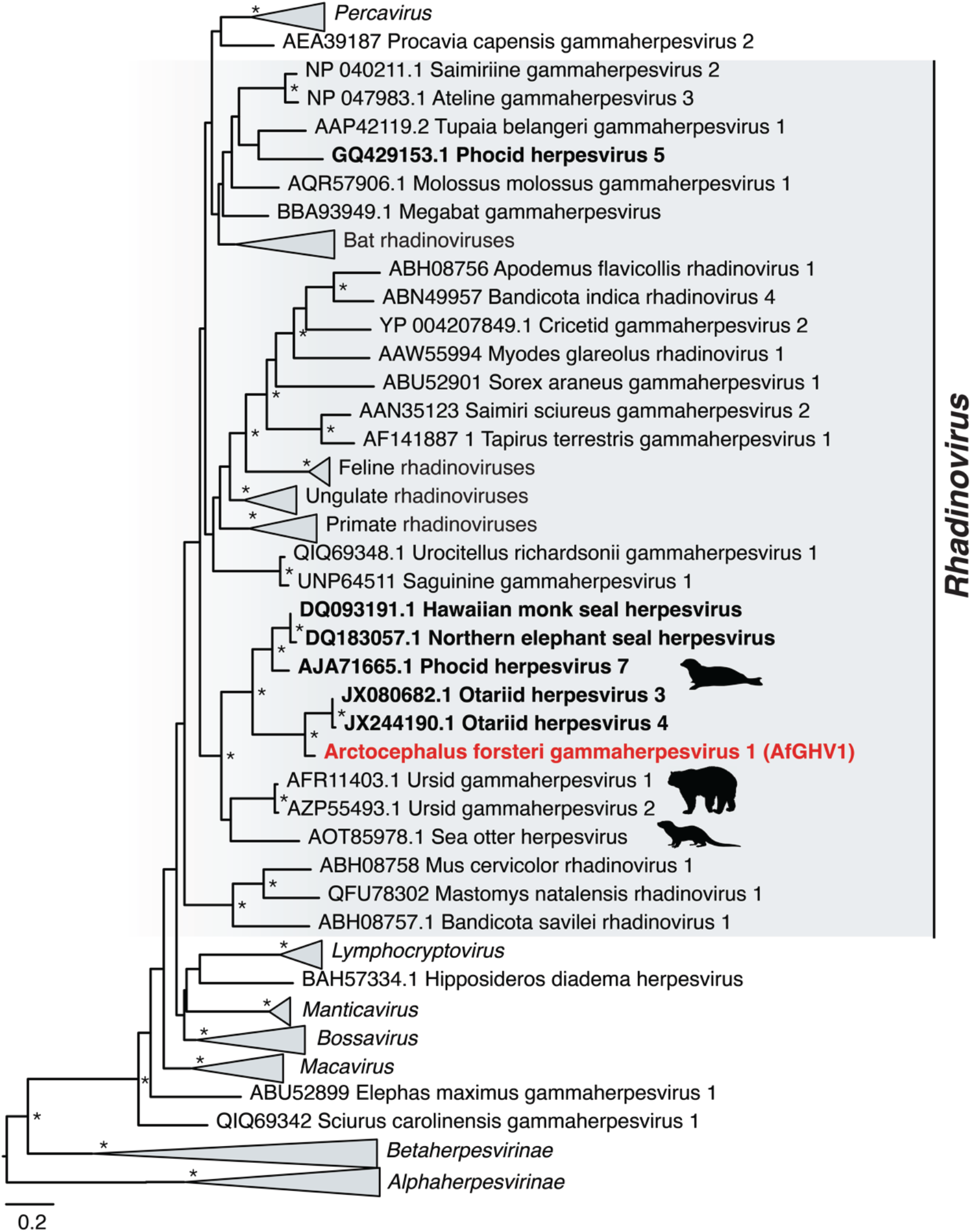
Phylogenetic relationships of the *Herpesviridae*. An ML phylogenetic tree based on the conserved amino acid sequences of the DNA polymerase gene with Arctocephalus forsteri gammaherpesvirus 1 (AfGHV1) shown in red and in the context of its closest relatives. The tip labels of pinniped associated herpesviruses are bolded. Animal silhouettes depict the virus-host associations for the taupapillomaviruses. All branches are scaled to the number of amino acid substitutions (model LG+F+R9) per site, and the tree is midpoint rooted for clarity only. An asterisk indicates node support where SH-aLRT >= 80% and UFboot >= 95%.

### 3.4 SL16 library composition

To examine whether other pathogens were present in SL16, contigs assembled from rRNA-depleted reads were assessed for taxonomic associations using CCMetagen. SL16 was predominately (85%) comprised of contigs belonging to eared seals (family *Otariidae*), while 5% was attributed to other pinnipeds (e.g., walruses), and non-chordates including bacteria (<0.1%) (Supplementary Figure 2A). Among the non-chordate abundance, 52% were associated with bacteria from various families, and 48% were linked to eukaryotes, specifically fungi (43%), arthropods (2%), algae (2%), and diatoms (1%) (Supplementary Table 4, Supplementary Figure 2B). The fungi were primarily assigned to *Candida albicans* (32%).

Further analysis of the CCMetagen results employing BLASTn revealed that a contig with hits to the *Tannerellaceae* (reference sequence *Tannerella forsythia*, HG784150.1) shared 81% nucleotide similarity (e-value = 7e-93) with the tetratricopeptide repeat protein of *T. forsythia*. The contigs associated with this species were highly fragmented so we were unable to make a definitive judgement regarding the taxonomy of this bacterium. Additionally, a fragment associated with the *Enterobacteriaceae* (reference sequence *Shigella dysenteriae* EU855235.1) was found to be identical to a hypothetical gene found in *Escherichia coli* (e-value = 0.0, EFO55302.1). The remaining contigs associated with *Enterobacteriaceae* share the greatest sequence similarity with *E. coli* (e-value = 0.0, 95% nucleotide identity, CP017061).

## 4. Discussion

We used a metatranscriptomic approach to identify potential viral aetiological agents in New Zealand fur seals with and without gross and microscopic pathology. Using this approach on a variety of tissues, we were able to identify two novel DNA viruses and a likely aetiology for the oral papilloma-like lesions described.

Papillomaviruses have been found in a wide range of mammalian species, with 85 hosts identified to date (Van Doorslaer et al., 2017). Despite this, our understanding of papillomaviruses in seals and their associated pathology remains limited. We identified a novel papillomavirus, the first described in fur seals, and the first report of oral papilloma-like lesions in this host group. This finding is consistent with other taupapillomaviruses identified in oral lesions from dogs and cats (Dunowska et al., 2014; Munday et al., 2016). Although such lesions are often of limited clinical significance—as observed in this case study where the lesions self-resolved—there is potential for these plaques to progress into invasive squamous cell carcinomas (Munday et al., 2016).

A critical factor in papillomavirus-induced oncogenesis is the integration of the virus into the host genome and its impact on the expression of the viral oncogenes E6/E7. These genes are negatively regulated by E2, and integration, which is frequently observed in the HPV (Arias-Pulido et al., 2006) can disrupt the E2 gene, leading to upregulation of the E6/E7 oncogenes (Münger et al., 2004). We present preliminary evidence that AfPV1 may exist in both episomal and integrated forms within this individual. The gene expression profile of AfPV1 infection, particularly the high E2/E6 ratio, is typically indicative of episomal infection in HPV16 infection, although exceptions exist (Qiu et al., 2021). Conversely, the AfPV1 E2 gene appears truncated, and a chimeric contig with a predicted E2 breakpoint was assembled, suggesting that AfPV1 may also be present in an integrated form.

The integration of HPV has been shown to trigger substantial host genome alterations resulting in the complete loss of function (Schmitz et al., 2012) or increased expression in target genes surrounding virus integration sites (Ojesina et al., 2014). Analysis of the host-associated regions of the chimeric sequence suggests that AfPV1 integration occurred in the vicinity of genes P2RY2 and MRPL2. Of note, the P2RY2 receptor has been shown to modulate virus yield, calcium homeostasis, and cell motility in cytomegalovirus-infected cells (Chen et al., 2019). Further investigation is required to confirm the presence of this chimera, the integration of AfPV1, and its effect on host gene expression.

While establishing causality is challenging, as papillomaviruses often asymptomatically infect the skin, there are several key pieces of evidence indicating that AfPV1 was infecting the New Zealand fur seal and may be associated with disease in these animals, although further investigation is warranted. AfPV1 appears to be highly abundant within the metatranscriptome data, with SL16 transcripts representing 0.18% of reads in a pool of three individuals, two of which were PCR-negative for this virus. AfPV1 was limited to an individual with papilloma-like lesions, as determined by gross and histological examination prior to the molecular discovery of this virus. This, together with the construction of the entire genome of AfPV1 from metatranscriptomic data, makes it unlikely that this sequence represents an endogenous viral element (i.e, a viral sequence that was passed vertically in the germline of the host) but rather the presence of both integration and episomal AfPV1 replication.

There is no evidence to suggest that another pathogen was responsible for the oral lesions of seal Registry #3254. *C. albicans* was the only microorganism detected in relatively high abundance (0.05% of SL16 abundance in CCMetagen) and is a common oral fungus, acting most commonly as a commensal organism. Although *C. albicans* can be an opportunistic pathogen, it is not known to cause papillary lesions, and these organisms were not evident within multiple histological sections of the lesions (Dunn et al., 1984). Bacteria were also present in the SL16 library, such as *E. coli* and a sequence with homology to *Tannerella sp.*, although in the latter, the highly fragmented nature of the sequences assembled prevented our ability to taxonomically classify this species further. While *E. coli* and certain *Tannerella* species (e.g., *T. forsythia*) are known to be pathogenic, they are not known to cause oral lesions consistent with those described (Gomes et al., 2016; Holt and Ebersole, 2005).

A phylogenetic analysis provides additional evidence that AfPV1 is indeed associated with a seal host: it groups with the Carnivora-associated taupapillomaviruses in a clade with seal and polecat papillomaviruses, LwPV5 (MG571094) (Smeele et al., 2018) and MpPV1 (NC_022253) (Smits et al., 2013). Although there is some evidence of recombination and host switching within the *Papillomaviridae*, the known taupapillomaviruses appear to have generally co-diverged with their hosts (Van Doorslaer, 2013).

Our survey also revealed the partial genome of a novel *Gammaherpesvirus* in both the DNA and RNA components extracted from the oral mucosa of seal Registry #10741. Although this seal did not display any visible signs of viral infection, gammaherpesviruses have been linked to diseases in pinnipeds in the past (Dagleish et al., 2013). Consequently, this discovery warrants further investigation into the presence and impact of gammaherpesviruses in New Zealand fur seals.

The absence of viruses across many of our samples could be attributed to numerous causes. If low abundance viruses are present, it is possible that we did not have the sequencing depth to recover these over the host signal, that viruses are too divergent to detect using similarity-based methods, or that this simply could reflect the absence of infection entirely at the point of sampling. This is supported by the absence of evidence of viral disease in the histopathology reported from these seals.

## 5. Conclusions

Through a metatranscriptomic survey of tissue samples from New Zealand fur seals, we identified two novel DNA viruses. The identification of AfPV1 in a seal exhibiting oral papilloma-like lesions, and AfGHV1 in an individual showing no clinical symptoms of infection, emphasise the value of combining metagenomic sequencing with conventional gross and histological examination to discover novel wildlife pathogens. Such work may ultimately contribute to improved disease detection, control, and conservation efforts. Additional research is necessary to ascertain the clinical relevance of these viruses in New Zealand fur seals.

## Acknowledgements

We thank Dr. Wei Shan Chang for assistance with the genome organisation graph. We acknowledge the University of Sydney’s high-performance computing cluster Artemis for providing the computing resources used for this study. We acknowledge the Taronga Conservation Society Australia and New South Wales Department of Environment for their ongoing funding of the Australian Registry of Wildlife Health, and we thank the NSW National Parks and Wildlife Service, Organisation for the Rescue and Research of Cetaceans in Australia, and Taronga Wildlife Hospital for their ongoing efforts to rescue, rehabilitate, and care for injured and debilitated seals across NSW. We thank PhyloPic (https://www.phylopic.org) for the animal silhouettes used in Figure 3 and 4 under Public Domain Dedication.

## Ethics

These samples were collected under the auspices of the Taronga Conservation Society Australia’s Opportunistic Sample Policy (approval no. R22D34), and pursuant to NSW Office of Environment and Heritage-issued scientific licenses SL10469 and SL100104. Samples from live seals were collected by a veterinarian for diagnostic purposes under License to Rehabilitate Injured, Sick or Orphaned Protected Wildlife (no. MWL000100542) issued by the New South Wales (NSW) Department of the Environment. Samples from deceased, beach cast or euthanased seals, were collected during routine necropsy for disease surveillance in accordance with NSW National Parks and Wildlife Act 1974, section 132c, Scientific License number SL100104.

## Supplementary Figures

**Supplementary Figure 1.**
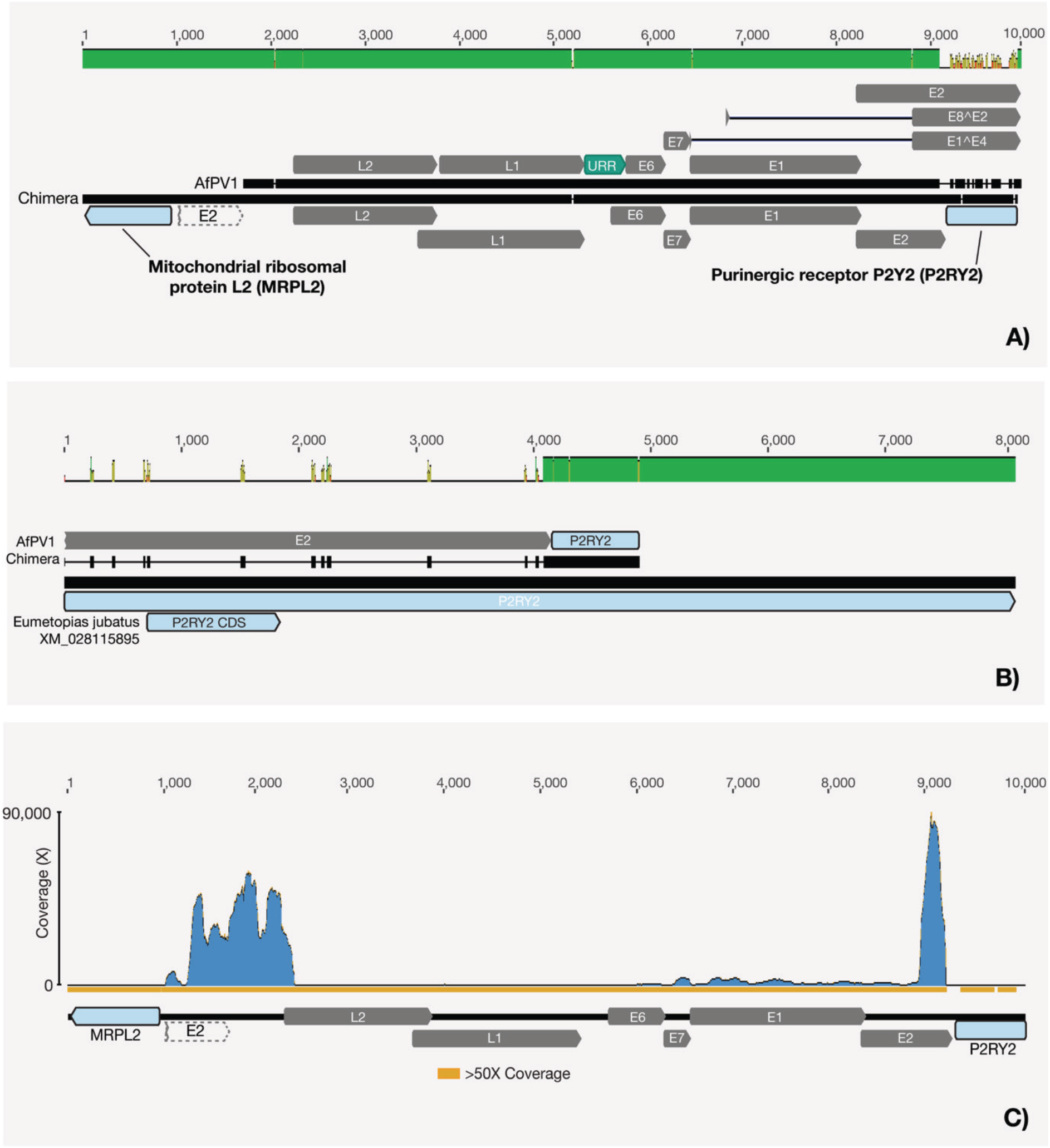
Genomic analysis of the Arctocephalus forsteri papillomavirus 1 (AfPV1) and the *Arctocephalus forsteri* chimera. (A) Nucleotide alignment comparing the genomic sequences of AfPV1 and an AfPV1-*Arctocephalus forsteri* chimera. The green bars situated above the alignment represent sequence identity, calculated with a window size of 1. The E2 annotation with dashed lines represents the region downstream from the breakpoint that no longer has a start codon. (B) Nucleotide alignment of the purinergic receptor P2Y2 fragment from the chimera and its closest relative in the northern sea lion (*Eumetopias jubatus*) XM_028115895. (C) Read coverage graph across the AfPV1 chimera. Regions with read coverage exceeding 50x are denoted by orange bars below the alignment, indicating areas of high sequencing depth. Across the panels, AfPV1 gene annotations are depicted in grey, while predicted host regions are shown in light blue.

**Supplementary Figure 2.**
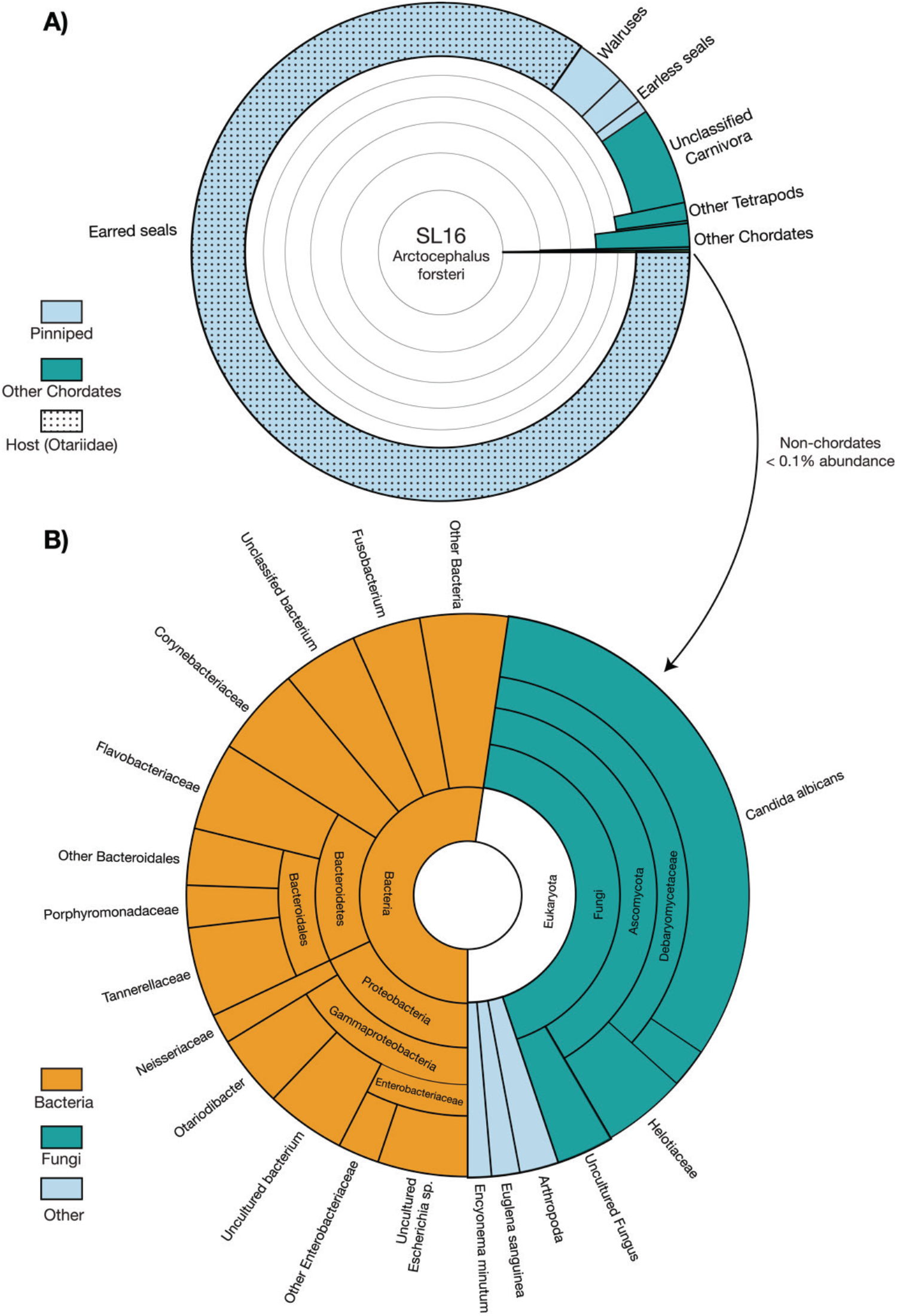
Taxonomic assignments of contigs in sequencing libraries. (A) Krona graphs illustrating the relative abundance of taxa in the SL16 metatranscriptome at varying taxonomic levels. For clarity, a maximum depth of six taxonomic levels was chosen. (B) Krona graph of the non-chordate subset of taxa. Across both panels segments are highlighted based on the species’ taxonomic grouping. Dots have been used to signify where contigs have been taxonomically assigned within the same family (*Otariidae*) as the New Zealand fur seal (*Arctocephalus forsteri*). Contigs without any matches in the database are not shown.

**Supplementary Table 1. Sample collection and library details**

**Supplementary Table 2. Predicted ORFs, protein products and binding sites**

**Supplementary Table 3. AfPV1 gene expression analysis**

**Supplementary Table 4. SL16 metatranscriptome composition**

