## Supplementary figures and images for "Identification of a novel papillomavirus from a New Zealand fur seal (*Arctocephalus forsteri*) with oral papilloma-like lesions"

### Supplementary Figure 1

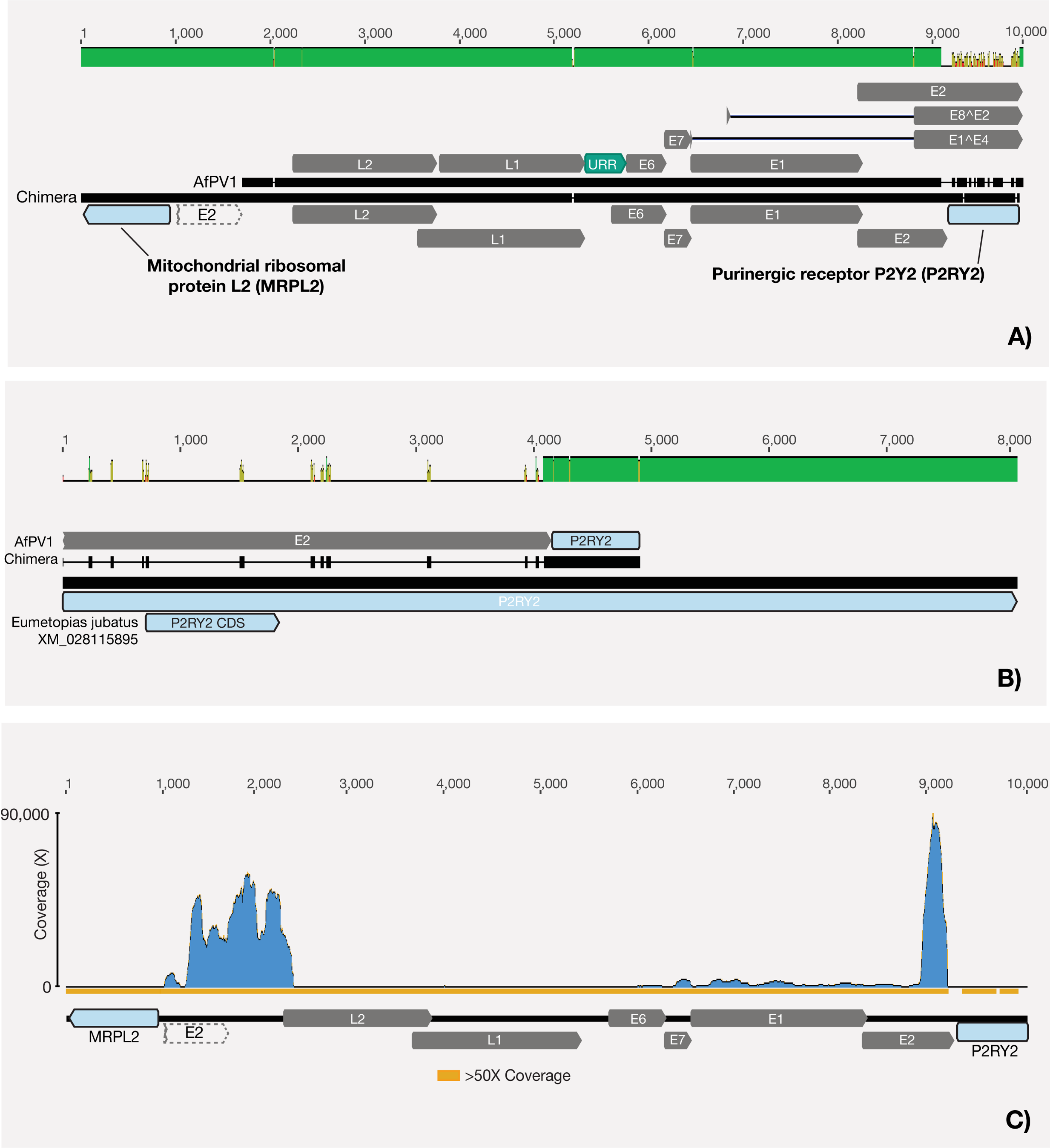

### Supplementary Figure 2

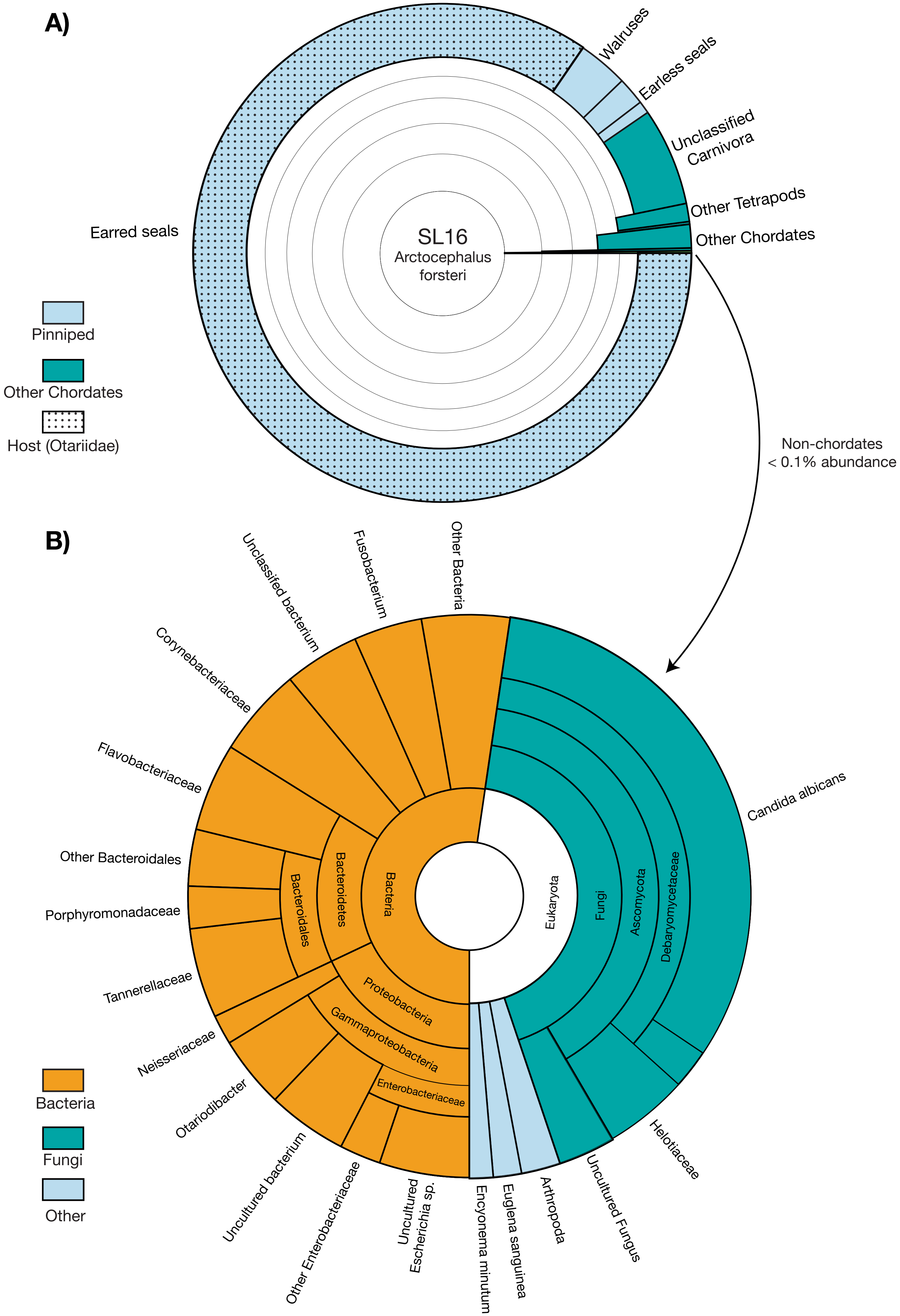
